# *Specifind*: A Natural Language Processing Tool for Automating Species Occurrence (Re-)Discovery from Scientific Literature

**DOI:** 10.64898/2025.12.24.696373

**Authors:** Golomb Durán Tomas, Far Antoni Josep, Díaz Anna, Barroso María, Roldán Alejandro, Colinas Noemi, Cancellario Tommaso

**Author notes:** **Corresponding author:** Golomb Durán Tomas **Mail:** CBB - Universitat de les Illes Balears (Spain).

## Abstract

A vast amount of valuable information on species occurrences remains embedded within the unstructured and continuously expanding body of ecological literature written in natural language. When effectively extracted, such dispersed knowledge has the potential to significantly improve understanding of species distributions and ecological patterns, thereby enabling more targeted and informed conservation actions. To address this challenge, Specifind is introduced as a tool designed to facilitate the extraction of species occurrence data through the identification of scientific species names, geographic references, and the relationships between them while ensuring traceability. At its core lies a newly developed and expertly annotated dataset comprising over one thousand open-access abstracts drawn from five domains: biogeography, botany, entomology, mycology, and zoology. A suite of Natural Language Processing components has been integrated, including Document Layout Analysis, Optical Character Recognition, Named Entity Recognition, Coreference Resolution, and Relation Extraction. These components enable the accurate identification and contextual linking of species and geographic entities, even when references are distributed throughout the text. As a result, Specifind enhances the discoverability and usability of species occurrence information embedded within unstructured scientific texts. It is expected to reduce the substantial effort required for manual literature review, thereby supporting biodiversity research, informing conservation planning, and facilitating ecological analysis.

## 1. Introduction

The spatial and temporal scales at which ecological processes operate often exceed those at which data are collected (Hampton et al., 2013). The understanding of species distributions and the prediction of responses to environmental change have historically been constrained by this mismatch (Franklin et al., 2017). Expansive biodiversity platforms are helping to bridge this gap by integrating observations from diverse sources into unified datasets spanning continents and centuries (König et al., 2019; Wüest et al., 2020). This platform-mediated integration represents a fundamental shift in ecological practice, moving the discipline toward increasingly computational and data-intensive approaches (Michener and Jones, 2012).

In the last years, platforms such as Global Biodiversity Information Facility (GBIF.org 2025), World Register of Marine Species (WoRMS Editorial Board 2025), iNaturalist (iNaturalist 2025), and Ocean Biodiversity Information System (OBIS 2025) have emerged as critical infrastructure for biodiversity research, aggregating billions of records from museum collections, field surveys, and citizen science initiatives worldwide. Despite their large utility, these centralized repositories still suffer from significant gaps of species coverage and the locations they inhabit (Yesson et al., 2007; Troia and McManamay, 2016; Hughes et al., 2021; De Araujo et al., 2022; Garcia-Rosello et al., 2023; Seebens et al., 2025). As a result, the lack of distribution data can potentially distort the view on large-scale biodiversity patterns (Beck et al., 2014).

Fortunately, much of this information to fill the gap on species knowledge exists, not in structured databases, but embedded within the prose of scientific publications (Mora et al., 2011; Thessen and Patterson, 2011). Research articles, including taxonomic descriptions, and technical documents like field reports and ecological surveys contain detailed information about where species have been observed, collected, or studied (Paterson et al., 2000). This textual knowledge spans centuries of fieldwork accumulating scientific discoveries that represent an invaluable and partially untapped resource for contemporary biodiversity research. Indeed, this potential has already begun to be realised, as existing literature has been exploited in a considerable number of ecological studies for the purpose of detecting trends and topics, evidence synthesis and literature reviews, expanding literature-based datasets, and extracting and integration of primary biodiversity data as reviewed by Farrell et al. (2022).

Traditionally, the extraction of this embedded knowledge has relied on manual curation by domain experts, who read publications and transcribe relevant information into databases. While this process may ensure high accuracy, it faces fundamental scalability limitations. The ecological literature grows exponentially (Cornford et al., 2020), with thousands of new articles published monthly across specialized journals. With the current publication rate, manual processing cannot match syntheses and compilation of literature-based datasets (Westgate et al., 2018), creating an ever-widening gap between available knowledge and accessible data. Therefore, developing informatics tools to automate the information extraction of meaningful, context-specific ecological information from unstructured scientific texts and transform it into structured data suitable for downstream analyses is crucial in the current ecological research.

One promising avenue for achieving this goal lies in Natural Language Processing (NLP), which enables automated extraction of structured information from text. Conventional approaches relied on lexicons for species name recognition, gazetteer matching for locations, and rule-based methods for associations (Gerner et al., 2010; Pafilis et al., 2013; Silva et al., 2016). While effective in constrained settings, these methods often failed to generalize across diverse writing styles, domain-specific vocabulary, complex sentence structures, and exhibit limited robustness when applied to previously unseen data.

Meanwhile, machine learning techniques, including Conditional Random Fields, Support Vector Machines, N-grams and Neural Networks improved flexibility but required extensive feature engineering and struggled to transfer across domains. The rise of deep learning further advanced the field, with Recurrent Neural Networks (RNN), particularly the Bi-directional Long Short Term Memory approach (BiLSTM), demonstrating superior performance on NLP tasks, as reported by Nguyen et al. (2019). BiLSTMs’ advantage stemmed from the ability to model contextual dependencies, reduce reliance on hand-crafted features, and learn richer semantic representations. However, RNN-based models, including BiLSTMs, still suffered from inherent sequential token processing constraints, limiting scalability in parallel computation, and difficulty capturing very long-range dependencies.

In recent years, multiple milestones in deep learning have transformed NLP. The revolution began with the introduction of the transformer architecture (Vaswani et al., 2017), which, through its self-attention mechanism, enables the modelling of complex contextual dependencies and parallel processing over long sequences. Building on this architecture, BERT (Devlin et al., 2019) demonstrated the power of large-scale pre-training using bidirectional transformers on extensive unlabelled corpora, popularizing the pre-training-fine-tuning paradigm and establishing it as the dominant approach in the field. In this paradigm, models are pre-trained in a self-supervised manner on unlabelled text and subsequently fine-tuned on downstream tasks through supervised learning. This approach allows knowledge to be transferred to domain-specific applications even when only limited labelled data are available (Min et al., 2021; Ding et al., 2023) and using minimal computational resources compared to training from scratch. In addition, Domain-Adaptive Pre-Training (DAPT) (Gururangan et al., 2020) emerged as an effective intermediate step between general purpose pre-training and fine-tuning. In DAPT, a pre-trained model is further trained on a large corpus of unlabelled domain-specific text, enabling it to better capture the syntactic and lexical patterns unique to the target field. Moreover, transformer-based models have become widely accessible through Hugging Face (Wolf et al., 2019), which provides open-source implementations, pre-trained weights, and easy-to-use API. This accessibility not only promotes reproducibility and transparency in research but also democratizes the use of advanced models (Riva et al., 2024). In the context of biodiversity informatics, these advances open new opportunities. Taxonomic name recognition has already benefited from the application of these innovations, as demonstrated by TaxoNERD (Le Guillarme and Thuiller, 2022), whilst the potential for extracting ecological interactions has been shown by Gabud et al. (2024).

Building upon these developments, Specifind is presented in this work as a newly developed tool designed for the processing of scientific texts. The system focuses on the identification of species names and geographic locations, as well as the relations between them that may constitute observations. The extracted information is subsequently converted into structured, machine-readable formats, while ensuring that traceability is maintained for each occurrence extracted. This approach aims to provide a scalable pathway for accessing species distribution information that was previously available only at considerable cost. The resulting data may be employed to fill existing spatial, temporal, and taxonomic gaps in biodiversity records, and to enable comprehensive legacy data integration. Ultimately, supporting more robust evaluations of species distributions and ecological dynamics.

## 2. Material and methods

A biological occurrence represents a core unit of biodiversity information, documenting the existence of an organism at a particular place (Darwin Core; Wieczorek et al., 2012). Whilst the Darwin Core standard includes a comprehensive set of metadata, Specifind focuses on extracting and georeferencing the spatial component of occurrence records from scientific literature. The following section describes data preparation, model training procedures, system architecture, and software usage.

### 2.1 Pipeline overview

Extracting species occurrences from scientific literature is an inherently complex task, due to the presence of multiple interdependent challenges. To manage the complexity, Specifind has been designed as a sequential pipeline, in which each stage addresses a specific subtask (see Figure 1 for a schematic representation of the pipeline).

**Figure 1.**
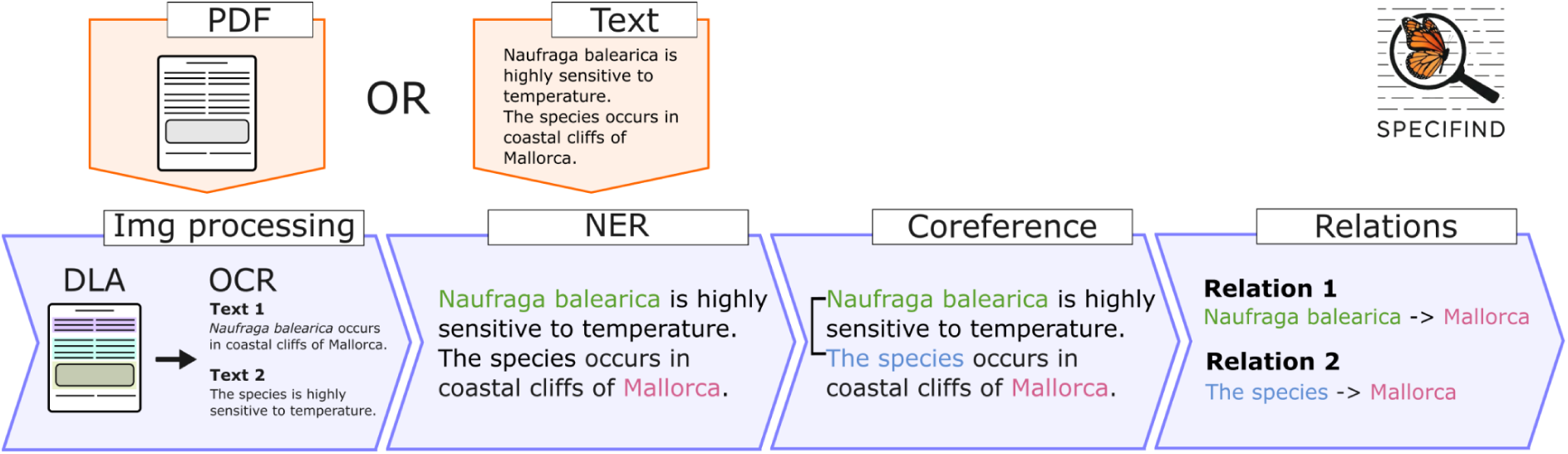
Schematic view of the main modules composing Specifind modules (purple) and the two entry points (orange), PDF or already available in raw text.

The predominance of scientific literature remains available exclusively in PDF format. This type of file is intended for visual presentation rather than for structured data storage. Once a document is saved as PDF, its textual content and underlying structure cannot be readily accessed, a phenomenon that has been termed as PDF prison (Agosti et al., 2019). To address this issue, Specifind incorporates a tailored component designed specifically for processing scientific publications to retrieve plain text and preserve document structure (Section 2.2).

Once textual content has been parsed, the next stage is Named Entity Recognition (NER), the purpose of which is to identify two types of entities: species names and geographic locations. This task is performed using a model fine-tuned on the dataset introduced in this study (Section 2.4). Following NER, Coreference Resolution (CR) is applied to associate indirect mentions with their corresponding entities, thereby enabling the full identification of organisms and locations throughout the text (Section 2.5). The pipeline proceeds with the Relation Extraction (RE) task for the identification of relationships between the two entity types that may indicate occurrences. The present dataset also includes annotations used to fine-tune a RE model as a text classifier (Section 2.6). The final output is a structured representation comprising a comprehensive list of identified species and geographic entities, along with the semantic relationships established between them. Each relation is linked to the original sentence in the text from which it was extracted, ensuring traceability and interpretability of the result, as well as enabling a faster and more efficient human validation.

To facilitate the efficient processing of textual data, all components were integrated into a spaCy pipeline (Honnibal et al., 2020) as its modular architecture allows seamless integration of custom text processing components within a unified environment. SpaCy is an open-source Python package that provides an object-oriented API for representing and handling linguistic data, together with robust and optimised tools, enabling high-performance processing of large corpora. Moreover, it is actively maintained and regularly updated to incorporate the latest advancements in natural language processing, ensuring compatibility with state-of-the-art NLP models and techniques.

### 2.2 Document text extraction

Scientific publications often exhibit complex and heterogeneous layouts, combining multi-column text, figures, tables, captions, and diverse typographic conventions. Such structural complexity poses significant challenges for automated document processing systems. Optical Character Recognition (OCR) is defined as the automated process of converting images, including scanned documents, photographs, or other visual representations of printed or handwritten text, into machine-encoded text. Conventional OCR systems are generally proficient at recognising individual characters and words; however, they do not inherently capture the overall structural organisation or the semantic relationships between distinct textual regions. Consequently, outputs generated by standard OCR pipelines are frequently fragmented or disordered.

To overcome these limitations and achieve coherent text reconstruction, Specifind combines OCR with the drive of Document Layout Analysis (DLA), which aids in the segmentation and interpretation of the spatial organisation of content on a page. Text extraction in Specifind is built upon the Surya (Paruchuri and Datalab Team, 2025) OCR and DLA fine-tuned vision transformers models supporting 93 languages.

The extraction pipeline operates in two stages. First, each page of the document is transformed into an image, after which DLA is applied to identify structural elements, their bounding boxes, and the reading order, followed by OCR. Only layout blocks corresponding to natural language content are retained (blocks labelled as Text, Caption, List Item, Section Header, and Form). Visual elements such as images, tables, and figures are excluded, as they convey information through visual rather than linguistic means.

Second, each detected OCR line is assigned to the layout block with which it shares the greatest bounding box overlap. Within each layout block, text lines are subsequently ordered by vertical position to preserve the original reading order. The final document text is reconstructed by concatenating lines within blocks using spaces and blocks themselves using line breaks, whilst applying dehyphenation to words split across lines.

### 2.3 Corpus and Annotation

#### 2.3.1 Data Collection

A novel, domain-specific, manually annotated corpus was constructed to support the training and evaluation of Specifind. The design of this resource was inspired by the COPIOUS dataset (Nguyen et al., 2019). The corpus is composed of abstracts, including titles, from scientific publications downloaded from the Scopus database, with a random 1% sample selected from each of five ecological and biological domains: biogeography, botany, entomology, mycology, and zoology (see *Supplementary Material 1 - S1* for the queries used). The inclusion of multiple ecological and biological subdomains aimed at increasing the generalisability of the system across related disciplines.

Abstracts were selected in preference to full-text articles for both practical and methodological reasons. Abstracts are typically more concise whilst containing key findings and frequently occurring entities such as species names and geographic locations, which are central to the targeted information extraction task. Moreover, their reduced length facilitates both manual annotation and computational processing, enabling the creation of a larger and more diverse dataset within manageable resource constraints. This strategy was intended to maximise linguistic diversity and contextual variation whilst minimising redundancy and annotation overhead.

Even the use of abstracts for fine-tuning does not preclude application to full-text documents due to the nature of transformer-based language models. Pre-trained transformers acquire linguistic representations from extensive corpora that typically include full-text documents, enabling generalisation beyond the specific text types used during fine-tuning (Beltagy et al., 2019; Devlin et al., 2019). Furthermore, entity and relation recognition are predominantly determined by local syntactic context within the model’s attention window rather than global document structure. Consequently, models fine-tuned on abstracts retain the capacity to process full-text documents through sliding-windows that preserve local contextual information.

#### 2.3.2 Data Labelling

All texts in the dataset were manually annotated to enable supervised training and evaluation. Annotation was conducted using Label Studio (Tkachenko et al., 2022), an open-source data-labelling platform that provides a customisable tagging interface and supports collaborative workflows.

The labelling procedure was undertaken by a team of annotators with academic backgrounds in biology or related life sciences, thereby ensuring appropriate domain expertise and familiarity with ecological conventions. To promote consistency and minimise inter-annotator variability, a comprehensive set of annotation guidelines was prepared in advance of each annotation phase. The dataset was partitioned amongst annotators without overlap, with each annotator responsible for a distinct subset of documents. Throughout the annotation process, annotators maintained regular communication to discuss ambiguous cases and edge scenarios. Such cases were resolved through collective discussion and documented as clarifications to the guidelines. Further details of the annotation guidelines are provided in the corresponding following subsections.

### 2.4 Named Entity Recognition

#### 2.4.1 Entity definition

Entities classified as species were annotated with a focus on taxonomic accuracy and biological specificity. Vernacular names and vague or generic taxonomic references (e.g., “birds,” “trees,” “insects”) and other taxonomic levels were excluded from annotation to preserve the precision and utility of the dataset as species level information is considered the most precise for occurrence data.

Geographical entities were annotated when they referred to real, identifiable spatial units. When a more precise location was provided, the entire entity was annotated. For instance, if both a city and its corresponding country or region were mentioned (e.g., “Mallorca, Balearic Islands, Spain”), the complete list of locations was annotated as a single geographic entity. This approach was intended to preserve contextual detail and enhance downstream utility for spatial analysis.

Detailed annotation rules for species names and geographical entities can be found in *Supplementary Material 1 - S2 and S3*.

#### 2.4.2 Model Training

The identification of species and location entities was undertaken through the fine-tuning of pre-trained transformer-based language models. This was implemented using a supervised training strategy and the domain-specific dataset, randomly partitioned into training (70%), validation (15%), and test (15%). An experiment was subsequently conducted using different BERT-based models to identify the model with the best performance (Tab. 1). Which, despite sharing a core transformer structure, differ in their training methodologies and data sources.

**Table 1:**
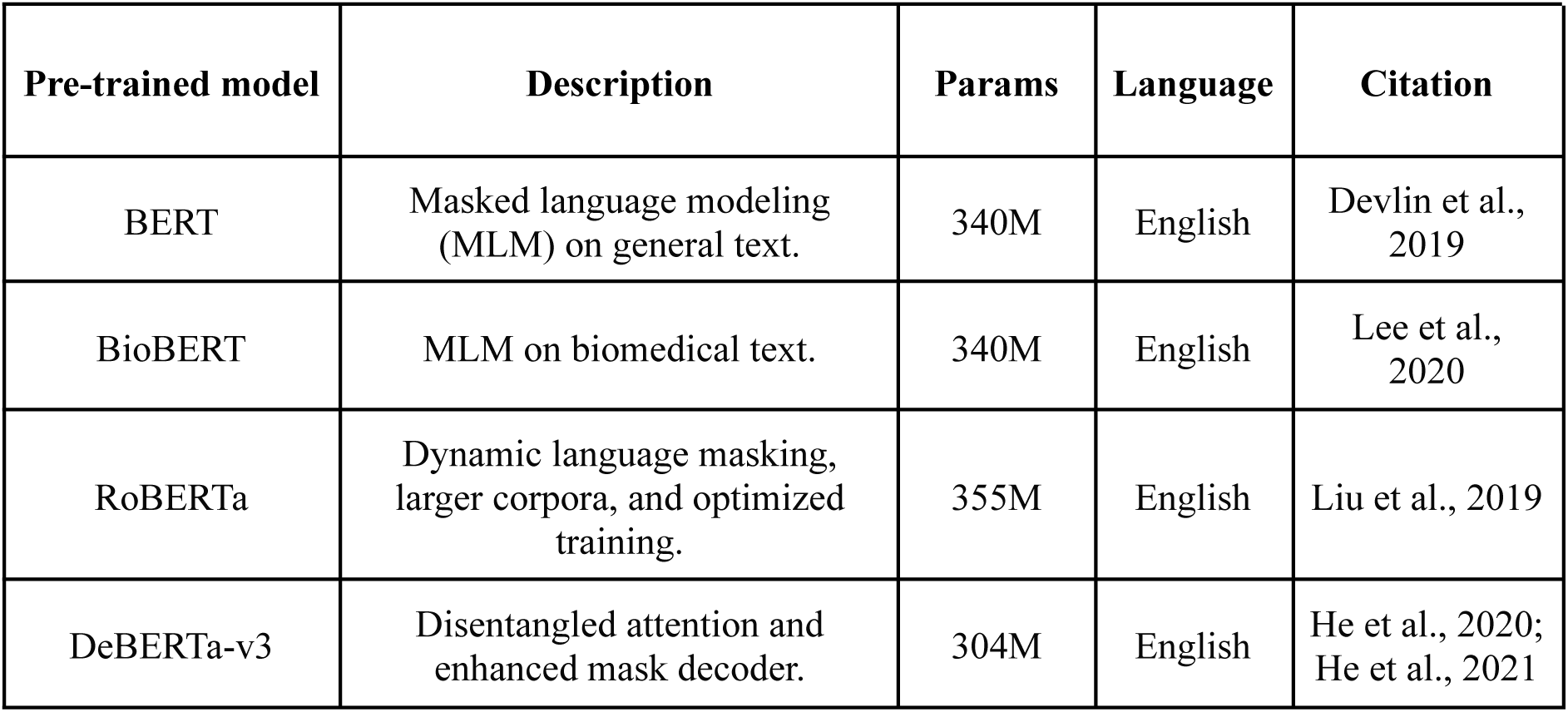
Overview of the BERT-based pre-trained language models used in the experiment, including their training corpus, architecture, and number of parameters (params).

To ensure a fair comparison and to maintain a controlled and reproducible environment for evaluating different transformer families, all NER models were trained using spaCy’s training framework. Employing a pipeline composed of a transformer component followed by a maxout architecture NER classifier with a hidden width of 64 and mean pooling over the transformer outputs. Cased and large size variants were selected; consequently, although SciBERT (Beltagy et al., 2019) was pre-trained on partially biomedical corpora, it was excluded as no large model size variant is available. Tokenization was performed using spaCy’s tokenizer and a maximum sequence length of 512 configured with strided spans (window 384, stride 128) to prevent truncation and properly handle texts larger than context window and mixed precision enabled.

Training was conducted on a RTX A6000 48gb VRAM GPU with a batch size of 128, gradient accumulation every 3 steps, a dropout rate of 0.1, and a maximum of 30 epochs. Optimisation followed the Adam algorithm with weight decay (L2 = 0.01), gradient clipping, and a warmup-linear learning rate schedule (initial rate 2×10⁻⁵, 250 warmup steps). Model selection was based on the F1 score of predicted entities, evaluated every 200 steps. Seeds were fixed to 42 for determinism.

### 2.5 Coreference Resolution

In natural language, coreference occurs when multiple terms refer to the same entity, known as the referent. It is a textual cohesion mechanism to avoid unnecessary repetition and contributes to the overall fluency of the text. These expressions often take the form of pronouns (e.g., “it”, “they”), definite nouns (e.g., “the species”, “this mammal”, “stripped dolphins”, “the study plane”), or other expressions referring to the full entity name. In NLP, Coreference Resolution (CR) is a well-established task aimed at identifying and grouping all textual mentions that refer to the same entity (Liu et al., 2023).

Specifind relies on the CR step to recover indirect references and linking them back to a precise species name or a geographic location. Anchoring mentions to one or more consistent representatives underpins the decision to exclude vernacular, vague or generic names during the NER scope (Section 2.4.1). To enable this feature, the pipeline integrates FastCoref (Otmazgin et al., 2022), a transformer-based model optimised for fast predictions with high-accuracy of various types of coreference, including pronominal, nominal, and demonstrative coreferences.

The predictions generated by FastCoref are organised into clusters, then Specifind identifies representative mentions for each cluster. These representatives are determined through the application of a relative minimum threshold based on the frequency of the NER entities within the cluster. This approach ensures that the most recurrent entities are chosen, including the detection of multiple head entities when the referent is plural (e.g. “***Podarcis pityusensis*** and ***Upupa epops*** are incredible animals. **These species** are present in the Balearic Islands.”) while minimizing the risk of selecting infrequent or incorrect predictions in large clusters.

### 2.6 Relationship Extraction

The species-location associations are derived from all possible pairs of the two entity types. Determining which candidate pairs correspond to valid associations constitutes the task of Relationship Extraction (RE) based on relative context. Specifind adopted a sentence-level approach rather than processing full texts. Accordingly, candidate pairs were restricted to sentences containing at least one mention of each entity type. This design choice was motivated by two primary considerations: First, the local context principle suggests that relationships between entities are typically expressed within proximate textual boundaries (Wang et al., 2020). Transformer-based models operate with context windows that are generally constrained to shorter text segments, making sentence-level granularity well-aligned with their architectural capabilities. By processing relationships at the sentence level, the most relevant contextual information surrounding entity mentions is captured whilst avoiding the inclusion of potentially confusing or irrelevant information from distant passages; Second, transformer-based architectures exhibit quadratic computational complexity with respect to input length (Vaswani et al., 2017). Consequently, sentence-level inputs offer substantially greater computational efficiency compared to full-text processing.

#### 2.6.1 Relationship definition

A secondary annotation process was performed by identifying all candidate sentences within the previously marked NER dataset using the sentence segmentation component of spaCy’s *en_core_web_trf* pipeline. Sentences were selected if they contained at least one of each entity type, a species name and a geographic location. Subsequently, each sentence was subjected to manual review to determine whether it expressed a meaningful species-location relationship following annotation rules described in *Supplementary Material 1 - S4*.

#### 2.6.2 Model Training

Prior training, a preprocessing step was performed whereby each annotated entity was enclosed using special markers to facilitate entity disambiguation by the model. Specifically, species mentions were encapsulated with the tags *[S]* and *[/S]* and geographic mentions with *[G]* and *[/G]* (e.g., “In open scrublands of [G] Menorca [/G], observations confirmed the presence of the [S] *Upupa epops* [/S].”). This approach has been shown to enhance the model’s ability to focus on relevant entities and their contextual interactions, thereby improving relation extraction performance (Soares et al., 2019; Zhou and Chen, 2021). The annotated dataset was then partitioned into training (70%), validation (15%), and test (15%) subsets using a stratified sampling strategy to mitigate bias arising from the tendency of sentences containing both species and geographic entities to be positive relations.

Following dataset preparation, a binary classification model was fine-tuned using HuggingFace‘s transformers training API on multiple pre-trained models similar to NER training in Section 2.4. Text inputs were subsequently tokenised with fixed-length padding and truncation adjusted to 512 tokens. Training was conducted for 10 epochs with a learning rate of 2×10⁻⁵, weight decay regularisation, and batch sizes of 16 for training, respectively. Model performance was assessed using the F1 score, owing to its ability to provide a balanced assessment for an imbalance classification task of both precision and recall in binary classification tasks. Seeds were fixed to 42 for determinism.

#### 2.6.3 Handling coreferences

To address potential limitations of sentence-level RE and capture relationships that span sentence boundaries, prior to relationship extraction model inference, coreference chains are identified and resolved, with each coreferent mention replaced by its corresponding NER entity. This enables the system to recognize when pronouns or alternative references in one sentence refer to entities mentioned elsewhere in the document, effectively extending the scope of relationship extraction to the document level whilst maintaining sentence-level processing efficiency.

For example, in the text “*Aspergillus fumigatus* was isolated from soil. This fungus is also found in Formentera”. The nominal coreference “This fungus” is replaced by the head entity “*Aspergillus fumigatus*”. Therefore, the model will receive the input “[S] *Aspergillus fumigatus* [/S] is also found in [G] Formentera [/G]”.

### 2.7 Usage

Specifind was designed with accessibility and straightforward integration into existing workflows. All dependencies are bundled within the package, eliminating the need for external installations. Getting started is as simple as running the following command in a Python environment: pip install specifind Interaction with the system is centred around a single class, Specifind, which handles models loading and subsequent analysis. The architecture of the pipeline enables a single instance to handle multiple texts without requiring reinitialization. Allowing to reduce computational overhead by maintaining one instance throughout a session.

The class provides two principal methods. The first, analyze, enables the direct processing of text strings. Options are available to activate coreference resolution (coref), while results can be returned either as structured outputs or as a spacy.Doc object (return_doc) to allow further downstream natural language processing.

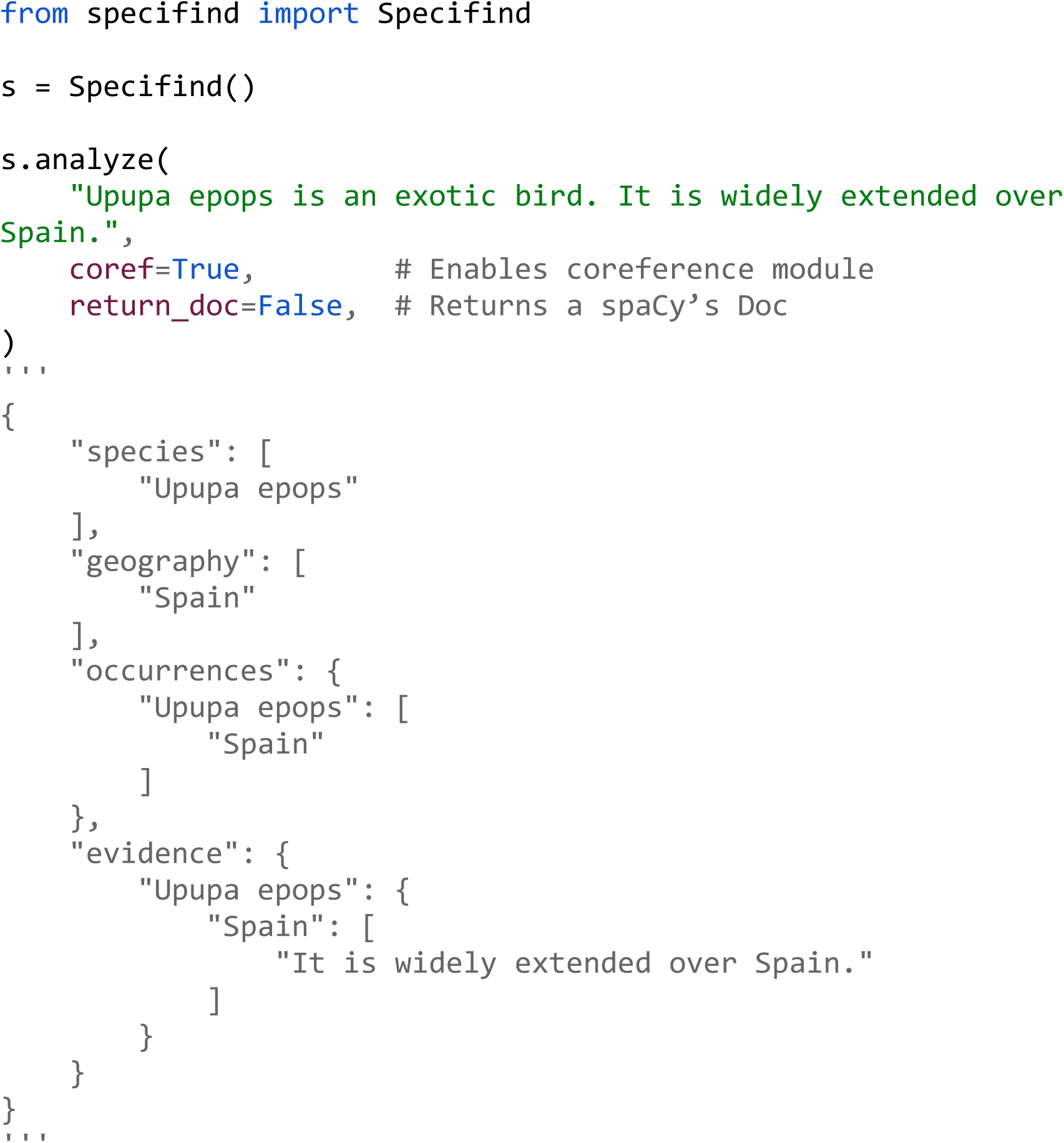

The second method, analyze_file, extends the system’s capabilities to include the parsing of PDF files through the integrated text extraction module. Additional parameters are available to allow the specification of page ranges (first_page, last_page), and the image resolution used during PDF page rasterisation (dpi). An increase in the dpi parameter results in higher-quality page images and, consequently, improved OCR accuracy; however, this improvement is accompanied by increased VRAM requirements for the underlying models when inferring on a GPU, which may lead to out-of-memory (OOM) exceptions in constrained environments.

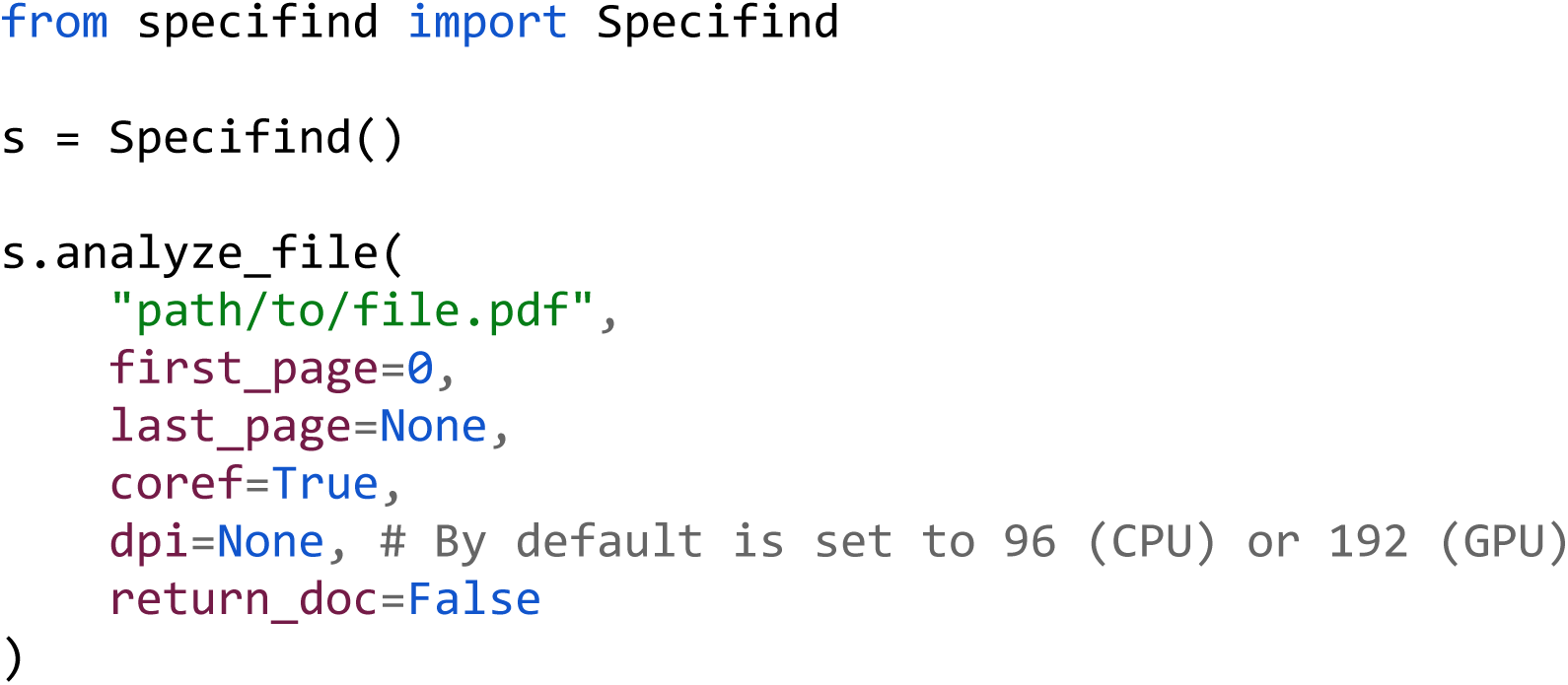

## 3. Results

### 3.1 Dataset

A total of 1,438 abstracts with a total count of 335,732 words. The collection included texts of varying length, with an average of 233 words (sd ± 87).

For the NER task, annotators recognized 3,800 species and 2,352 locations across the corpus. Of these, 1,925 unique species names and 1,283 unique location entities were distinguished (Fig. 2).

**Figure 2.**
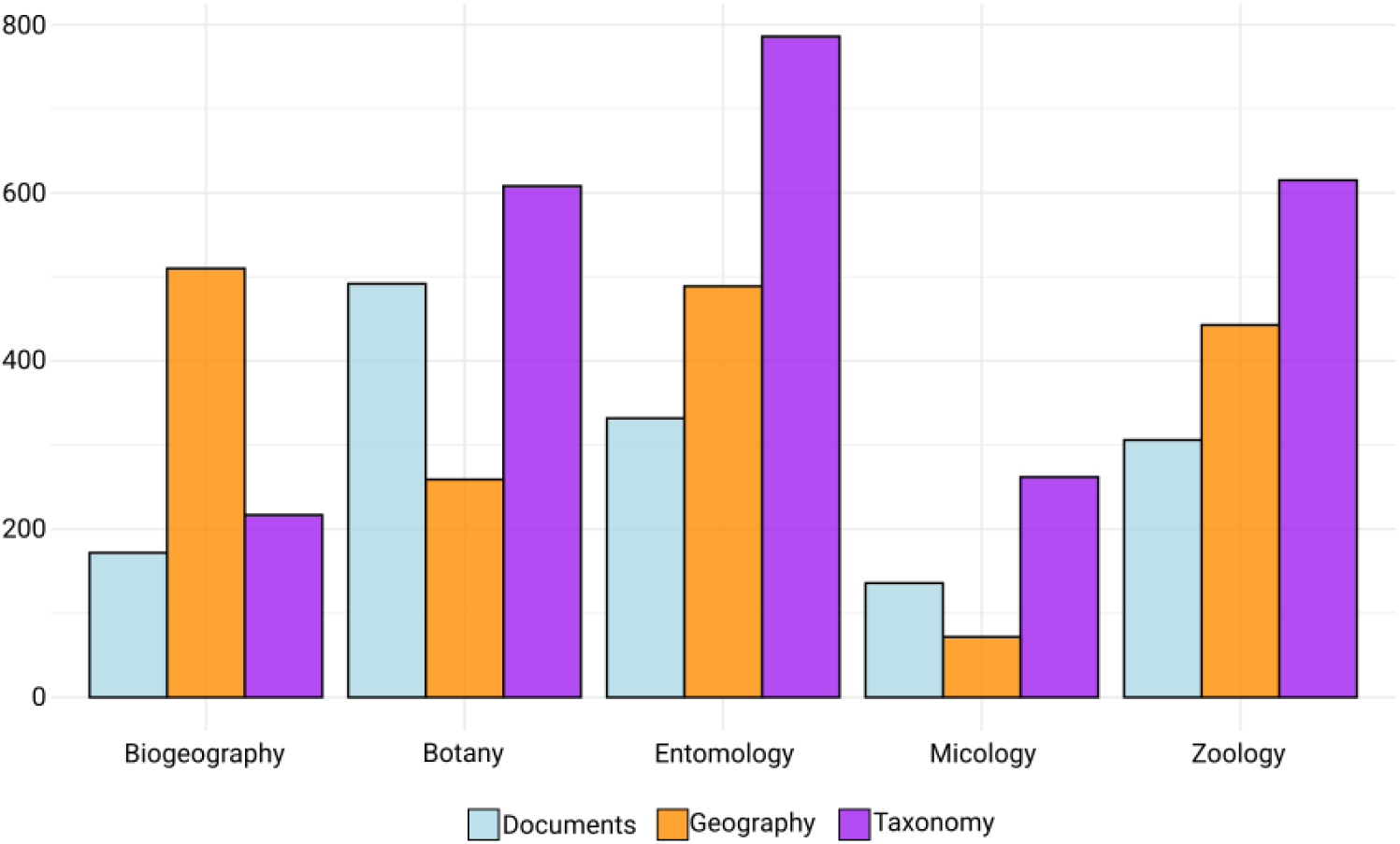
Count of entity type and number of documents per domain. Lightblue: document; Orange: Geography: Purple: Taxonomy.

Concerning the relationship extraction, a total of 1,179 candidate species-location associations were identified. Following the annotation process, 948 were confirmed as positive relations, while 231 were determined to be non-related (Fig. 3).

**Figure 3.**
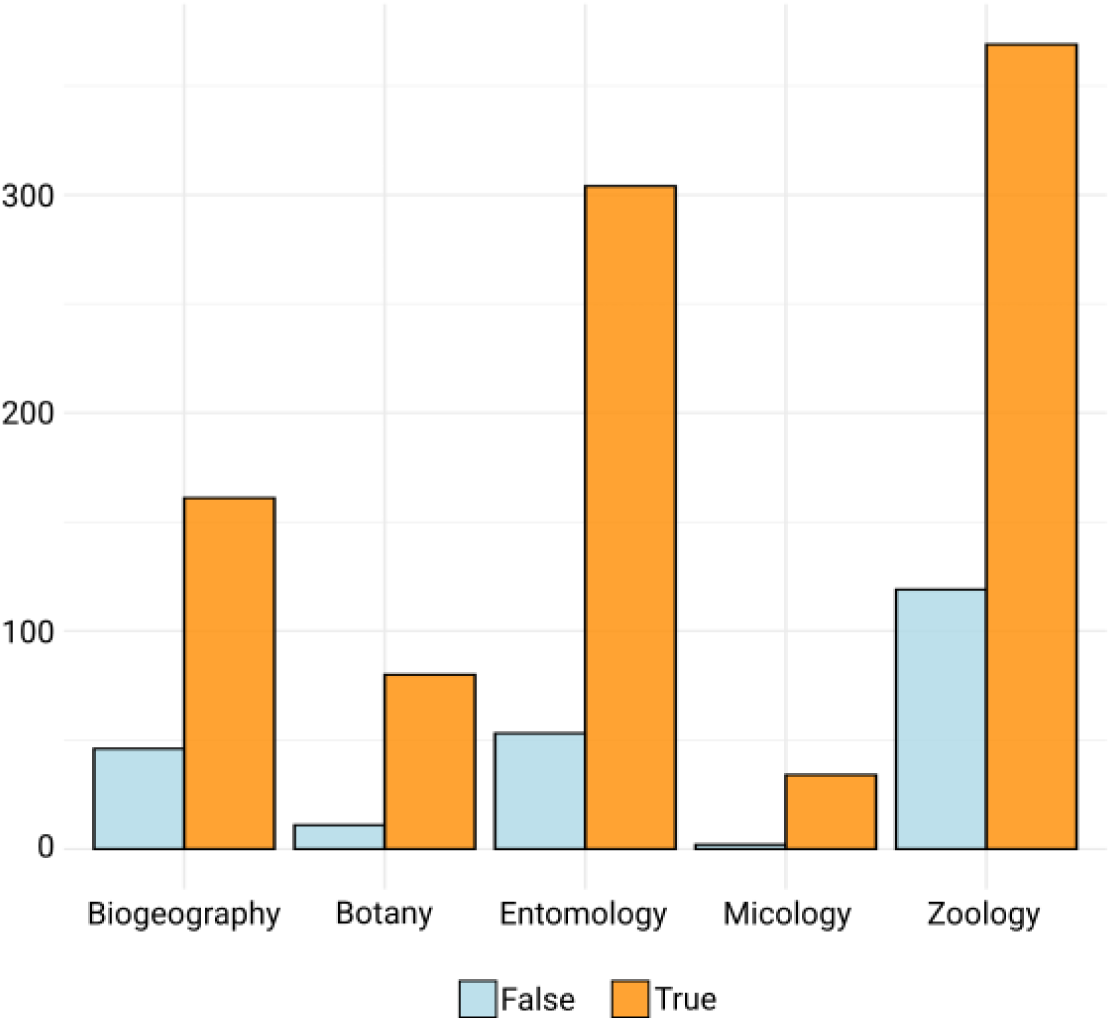
Count of positive (orange) and negative (lightblue) relationships between entities for each domain.

### 3.2 Named Entity Recognition

Precision, recall and F1 score for each of the models run on the experiment are presented in Table 2. The findings support the benefits of DAPT in performance, as evidenced by the species name with the highest F1 score achieved by BioBERT, representing an improvement of +2.1% relative to BERT. DeBERTa-v3 yielded the best performance, obtaining the highest F1 score across entities and supporting the effectiveness of its disentangled attention mechanism and enhanced mask decoder, all with a shorter parameter count.

**Table 2:**
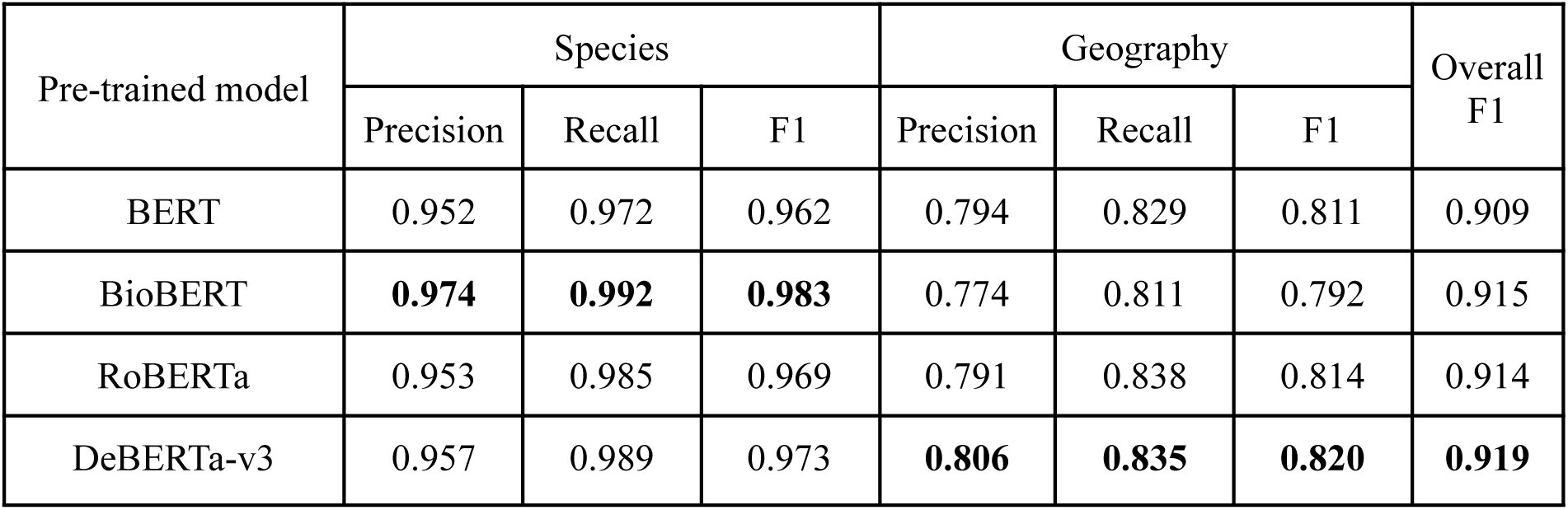
Benchmark results for different pre-trained models and fine-tuned over the same dataset. Best metrics between models are marked in bold.

Manual review of the predictions of geographic entities revealed a consistent shortcoming across models in accurately capturing long geographic chains. For example, annotated spans such as *“*western coastal region of Mallorca, Spain*”* were frequently truncated by the models to *“*Mallorca, Spain*”*.

To produce a more realistic evaluation of model performance, a partial matching heuristic evaluation was performed. Predicted entities were considered valid if they intersected with any part of the annotated location span, acknowledging partial overlap as an acceptable match in cases where the full chain was not captured. The results presented in Table 3 indicate an improvement of +10.6% in F1 score for geography entity recognition when comparing partial and exact matches, suggesting a tendency for large spans to be truncated.

**Table 3:** Comparison between exact match and partial match evaluation strategies for geography entities on the fine-tuned DeBERTa-v3-large. Best metrics between models are marked in bold.

| Match strategy | Species |  |  | Geography |  |  | Overall F1 |
| --- | --- | --- | --- | --- | --- | --- | --- |
|  | Precision | Recall | F1 | Precision | Recall | F1 |  |
| Exact | 0.957 | 0.989 | 0.973 | 0.806 | 0.835 | 0.820 | 0.919 |
| Partial/Intersect | **0.966** | **0.995** | **0.980** | **0.910** | **0.943** | **0.926** | **0.958** |

A more detailed examination of the incorrect cases revealed that many were not due to limitations of the model itself but rather arose from misannotated by human reviewers.

### 3.2 Coreference Resolution

The integration of CR produced a marked increase in the number of extracted species occurrences. The integration of FastCoref was assessed using the complete dataset, resulting in raising the count of occurrences from 948 of the original annotations as mentioned in Section 3.1, to 1,196, representing a 26.16% increase when CR is applied.

In addition to improving occurrence recovery, the use of CR allowed indirect mentions to be handled outside the NER stage, therefore the domain becomes more constrained. This separation reduced the complexity, although not formally quantified here, this simplification is likely to translate into more consistent annotations and potentially more stable training behaviour.

### 3.3 Relationship Extraction

The benchmarks presented in Table 4 summarize the performance of several transformer-based models on the RE task. Similar to NER, multiple architectures were evaluated under the same controlled environment. All models demonstrated strong performance, with F1 scores consistently above 0.96. Among them, RoBERTa achieved the highest F1 score of 0.97, slightly outperforming the other models.

**Table 4:** Benchmarks of the fine-tuned relationship extraction model.

| Pre-trained model | Precision | Recall | F1 |
| --- | --- | --- | --- |
| BERT | 0.944 | 0.993 | 0.968 |
| BioBERT | 0.943 | 0.985 | 0.964 |
| RoBERTa | **0.964** | **0.993** | **0.978** |
| DeBERTa-v3 | 0.950 | 0.985 | 0.967 |

Closer inspection of the error cases revealed that a significant proportion did not stem from model shortcomings, but rather from annotation inconsistencies within the corpus. In particular, several false positives corresponded to valid species-location relations that had been missed or incorrectly labelled by human annotators.

## 4. Discussion

The present work has demonstrated the feasibility and potential of employing NLP techniques for the systematic extraction of species occurrence data from unstructured scientific literature. Through the integration of multiple NLP components within a unified, open-source pipeline, Specifind accurately identifies species names and associated geographic locations while preserving the traceability of extracted information. The system’s architecture ensures ease of installation and use, making advanced text mining capabilities accessible to researchers without requiring extensive computational expertise. To the best of the authors’ knowledge, this represents the first tool to combine these capabilities in an integrated biodiversity-focused framework, constituting a notable advancement over labour-intensive manual literature review and traditional informatic methodologies.

Early rule-based and hybrid systems for species name recognition, such as LINNAEUS (Gerner et al., 2010) and OrganismTagger (Naderi et al., 2011), established that scientific names could be reliably located and normalised in text, thereby inspiring a family of dedicated taxonomic name-finding tools. Machine learning techniques applied in COPIOUS (Nguyen et al., 2019) and BiodivNERE (Abdelmageed et al., 2022) achieved considerable progress in extracting species occurrences and other species-related metadata. More recently, TaxoNERD (Le Guillarme and Thuiller, 2022) and Gabud et al. (2024) introduced deep learning methods tailored to the biodiversity domain, showing that transformer-based architectures can substantially improve performance.

Specifind advances this lineage by combining multiple state-of-the-art components with newly trained domain-specific models to automate species occurrence extraction from scientific literature. These improvements are underpinned by the revolution of transformer architecture and the pre-training-fine-tuning paradigm, which together enable competitive performance even with comparatively small annotated datasets.

Integrating coreference resolution as a separate stage improves pipeline robustness by allowing NER and RE to operate on well-defined entity mentions, while coreference handles referential variation across text, reducing annotation ambiguity and error propagation. Prior work of Le Guillarme and Thuiller (2022), has shown that loosely constrained or domain-mismatched entity definitions can substantially degrade NER performance, supporting the benefits of deferring reference resolution to specialized components.

In addition, the system provides a comprehensive, user-friendly, ready-to-use, and easily installable solution for biodiversity-focused information extraction. Specifind includes a built-in text extraction component that addresses challenges associated with PDF processing (Agosti et al., 2019). The system operates across multiple devices and supports both CPU and GPU execution for enhanced computational efficiency. Furthermore, Specifind supports usage in both Python and R through the reticulate package (Ushey et al., 2025), enhancing accessibility for the broader biological research community, for which R remains a predominant programming language.

While overall system performance is strong, failures inherent to statistical models may still occur. Variations in text style, the presence of highly domain-specific terminology, and inherent natural language ambiguities such as common versus scientific names, homonyms of taxon terms, and overlapping place names can occasionally lead to misclassification or incomplete linkage. These limitations are mitigated through the traceability features of Specifind, which preserve detailed provenance and full document analysis, enabling users to validate results and ensure reproducibility.

The implications of these findings are substantial. By enabling rapid and accurate extraction of species occurrence information, Specifind has the potential to enhance ecological analyses, inform conservation prioritisation, and support evidence-based decision-making. Potential applications include augmenting biodiversity databases such as GBIF, WoRMS, and OBIS; rapid assessment of species distributions for IUCN Red List updates; and monitoring of spread of invasive species starting from literature. Automated extraction further enables large-scale analyses that would otherwise be infeasible due to the time constraints of manual curation, a critical advantage in the context of accelerating biodiversity loss and the urgent need for timely data to guide conservation strategies.

Several avenues for future research emerge from this work. To date, no DAPT has been carried out specifically for the ecological literature domain, with most efforts remaining focused on biomedical domains. Applying DAPT to large ecological corpora is likely to yield significant improvements in recognising and disambiguating domain-specific terminology. Additionally, Specifind has been observed in practice to operate across multiple languages, formal evidence of its multilingual generalisability has not yet been established. The development of a multilingual annotated dataset, combined with the fine-tuning on cross-lingual models, for example XLM-RoBERTa (Lample and Connea., 2019), would provide a rigorous evaluation. Moreover, advances of vision transformers such as Donut (Kim et al., 2022) and LayoutLM (Huang et al., 2022) demonstrate strong performance in extracting textual representations from images and tables. Incorporating such models would enable the system to process general visual materials by converting them into textual descriptions that can be fed directly into Specifind, supporting multimodal analysis without requiring modifications to its core architecture. Recent research has also explored the use of generative large language models for information extraction, relying on prompting strategies to produce structured outputs from unstructured text. This paradigm differs fundamentally from the modular, pipeline-based approach adopted in Specifind, as exemplified by the ARETE R package (Branco et al., 2025), which demonstrates the potential for retrieving species occurrence information.

## Data accessibility

To facilitate reproducibility, original pipeline code, latest releases, training dataset, evaluation scripts, and documentation are available on the associated GitHub repository (https://github.com/ToGo347/Specifind and https://github.com/ToGo347/science-ocr).

## Acknowledgements

This study has been partially funded by GOIB/Conselleria d’Educació i Universitats through the project “SINCO2022/18146” and co-funded by the European Union.

This work has been partially funded and promoted by the Comunitat Autonoma de les Illes Balears through the Conselleria d’Educació i Universitats and by the European Union-Next Generation EU/PRTR-C17. I1 (SINCO2022/6717). Nevertheless, the views and opinions expressed are solely those of the author or authors, and do not necessarily reflect those of the Conselleria d’Educació i Universitats, the European Union or the European Commission. Therefore, none of these organizations shall be held liable.

We thank María Capa and Enrique Arboleda for taking the time to read this manuscript and provide valuable comments. We also acknowledge Claire Newman for her assistance in labelling the training dataset.

## Authors’ contribution

GDT, and CT conceived the initial idea of the study. GDT developed the methodology, Specifind, and wrote the original manuscript. CT provided ecological expertise, assisted with methodology, data annotation, and contributed to write the manuscript. FAJ, RA, and CN contributed to software testing and manuscript review. DA and BM contributed to data annotation and manuscript review. All authors read and approved the final manuscript.

