## Supplementary 1 for "*Specifind*: A Natural Language Processing Tool for Automating Species Occurrence (Re-)Discovery from Scientific Literature"

**Article type**: Article
**Title:** *Specifind*: A Natural Language Processing Tool for Automating Species Occurrence (Re-)Discovery from Scientific Literature

**Authors:** Golomb Durán Tomas[1*], Far Antoni Josep[1], Díaz Anna[1], Barroso María[1], Roldán Alejandro[1], Colinas Noemi[1], Cancellario Tommaso[1]

**Affiliation:**
[1]Centre Balear de Biodiversitat, Universitat de les Illes Balears, Palma, Spain

*** Corresponding author:** Golomb Durán Tomas
**Mail:**
CBB - Universitat de les Illes Balears (Spain)

**ORCID:**
GT 0009-0003-0430-2073
FAJ 0009-0008-8869-6482
DA 0000-0002-9706-8954
BM 0000-0001-9624-3602
RA 0009-0009-5040-1009
CN 0000-0002-7196-4224
CT 0000-0002-6637-4764

**Supplementary Material 1**

**S1.** Scopus Queries for Dataset Download.

**S2.** Annotation Rules for Species Names.

**S3.** Annotation Rules for Geographical Entities.

**S4.** Annotation Rules for Relationships Between Species and Geography.

**S5.** Species name recognition using Specifind in R.

**S1.** Scopus Queries for Dataset Download.

This section provides the Scopus search queries used to compile the textual corpus for the study. All searches were performed using standardized Boolean operators, controlled vocabulary, and field-specific filters.

`TITLE-ABS-KEY(domain)`: retrieves publications that contain the domain in the title, abstract, or keywords.

`LIMIT-TO(OA, "all")`: restricts results to publications available as open access.

TITLE-ABS-KEY(botany) AND ( LIMIT-TO ( OA,"all" ) )

TITLE-ABS-KEY(biogeography) AND ( LIMIT-TO ( OA,"all" ) )

TITLE-ABS-KEY(entomology) AND ( LIMIT-TO ( OA,"all" ) )

TITLE-ABS-KEY(mycology) AND ( LIMIT-TO ( OA,"all" ) )

TITLE-ABS-KEY(zoology) AND ( LIMIT-TO ( OA,"all" ) )

**S2.** Annotation Rules for Species Names.

Species entities were defined as scientific taxonomic names referring to biological species. Inclusion and exclusion criteria are described below.

**Inclusion criteria**:

- Binomial names at species taxonomic level.

  *e.g., New data about distribution and biology of [Sharpia rubida].*

- Abbreviated genus names are retained exactly as they appear.

  *e.g., [C. haemorrhoidalis]*

- Subgenus designations, when present (e.g., *Aedes (Stegomyia) albopictus*), must be included in the annotated span.

  *e.g., [Aedes (Stegomyia) albopictus]*

**Exclusion criteria:**

- Abbreviated species epithet due to ambiguity and lack of standardization.

  *e.g., C. v. viridis*

- Vernacular names.

  *e.g., bird, tree, bat*

- Authorship information must not be included in the annotated span.

  *e.g., [Pollentia perezi] Capa, Pons & Jaume, 2022*

- Modifiers or contextual descriptors must not be included in the annotation span.

  *e.g., [Aeromonas hydrophila]-infected*

**S3.** Annotation Rules for Geographical Entities.

Geographic entities were defined as explicit references to spatial locations, including countries, regions, administrative units, and other locality names. Inclusion and exclusion criteria are described below.

**Inclusion criteria**:

- Geopolitical units: countries, states, provinces, cities, towns or similar.
  *e.g., [USA], [Rio Branco], [Illes Balears]*
- Physical and natural regions: mountains, islands, rivers, lakes, ocean basins.
  *e.g., [Mount Malindang], [Balayan Bay], [Mediterranean Basin]*
- Conservation or administrative areas: national parks, reserves, protected zones.
  *e.g., [Chukotka (Western Beringia)], [Municipal Health Department in Rio Branco city]*
- Geographic coordinates (including both numeric and textual elements).
  *e.g., [13° 21' 30" N., 120° 30' 33" E.], [East coast of Mindoro]*
- Historical regions: extinct geopolitical entities or paleogeographic regions.
  *e.g., [URSS], [Pangea]*
- Full location chains when available: annotate the entire phrase when it increases precision.
  *e.g., [Mallorca, Illes Balears, España], [Western France]*
- Abbreviations of place names.
  *e.g., [UK], [USA]*

**Exclusion criteria**:

- Adjectival uses of place names.

  *e.g., "[Philippine] Government", "[Brazilian] people"*

- Institution or company names where the location is part of the organization's name.

  *e.g., [University of Balearic Islands], [Informa UK Limited]*

- Cultural or archival references when not denoting physical locations.

  *e.g., "Archival fond of the National Academy of Sciences of Ukraine"*

- Ambiguous references where coordinates cannot be reliably assigned as multiple places can be addressed.

  *e.g., [mountains of Greece], [forest regions of Gabon]*

- Ambiguous or incomplete coordinate references with multiple plausible matches.

  *e.g., unclear or vague coordinate strings without clear spatial disambiguation*

- Coordinate ranges.

  *e.g., 5° and 15° S*

- Coordinates not properly formatted (eg. Words in between).

  *e.g., verbatimLatitude: 36°20'60"S; verbatimLongitude: 73°43'60"W*

**S4.** Annotation Rules for Relationships Between Species and Geography.

Annotations focused on explicit associations linking taxonomic geographic entities. Relationships were annotated only when supported by clear contextual evidence in the text, ensuring consistency annotations.

- **Related**: The sentence indicates that the species had been observed, found, sampled, distributed, or otherwise associated with the location in question, regardless of whether it is or was considered extinct.
- **Non-related**: Species and location mentions co-occurred without a clearly expressed or interpretable relationship, such as those explicitly indicating the species was absent, or otherwise not affirming any association.

**S5.** Species name recognition using Specifind in R.

Specifind was accessed from R using the *reticulate* (Ushey, Allaire & Tang 2025) package to enable easy integration between R and Python languages. To use Specifind requires a Python version greater than 3.10 (here Python 3.11 was used). Specifind and its dependencies were installed within a dedicated Python virtual environment, which was explicitly configured to ensure computational reproducibility and isolation from system-level libraries.

Once the virtual environment was activated and linked to R via *reticulate*, the *Specifind* class was imported and instantiated directly within the R session. The 'analyze' function was then applied to textual input to extract taxonomic and geographic entities, as well as to infer their contextual relationships within the text. Coreference resolution was enabled to improve entity linking across sentences. The output consisted of structured data describing detected species mentions, associated locations, and relevant contextual attributes.

All analyses were performed on macOS.

1) Open the terminal and install Python 3.11, create a virtual environment.

```
# /opt/homebrew/bin/python3.11 -m venv ~/specifind-env

# source ~/specifind-env/bin/activate
```

2) Install Specifind

```
# pip install --upgrade pip

# pip install numpy
git+https://github.com/ToGo347/Specifind.git
```

3) Open R, load the reticulate library and configure the virtual environment.

```
# Load reticulate package
```

```r
library(reticulate)

use_virtualenv("~/specifind-env", required = TRUE)

py_config(); py_list_packages()
```

4) Import Specifind and run the analysis.

```r
# Import the Specifind class

Specifind <- import("specifind")$Specifind


# Create an instance

s <- Specifind()


# Run the analysis

text <- "Upupa epops is an exotic bird. It is widely extended
over Spain."

result <- s$analyze(text, coref = TRUE, return_doc = NULL)
```
